## Supplemental Figures for "A framework for multiplex imaging optimization and reproducible analysis"

<u>List of Supplementary Figures and Tables</u>

Supplementary Figure 1: Standard IF versus CycIF in Normal Breast and HER2+ Tumor Tissue.
Supplementary Figure 2: Regional Variation in Threshold Results.
Supplementary Figure 3. Tissue Retention during CyCIF.
Supplementary Figure 4. 15-Minute Quenching in Different $H_2O_2$ Concentrations.
Supplementary Figure 5. Quenching under Different Time and Light Conditions.
Supplementary Figure 6: First versus second antibody application to same TMA tissue.
Supplementary Figure 7: Quantification of first versus second antibody application to same TMA tissue.
Supplementary Figure 8: Variables Impacting Round-Effect of Stain Quality and Bleed Through Observed in Miltenyi MACSima prototype Instrument.
Supplementary Figure 9: Single Cell Effect of Quenching in Pancreas Tissue.
Supplementary Figure 10: Single Cell Effect of Quenching in 72-core Tissue Microarray.
Supplementary Figure 11. Single Cell Annotation for Evaluation of Autofluorescence Subtraction.
Supplementary Figure 12. Single Cell Effect of Quenching in 11-core HER2+ Tumor Tissue Microarray.
Supplementary Figure 13: Replicates of Three HER2+ Tumor TMA Cores
Supplementary Figure 14: Comparison of Manual Versus Automated SBR
Supplementary Figure 15: kBET evaluation of batch correction methods in HER2+ breast cancer tissue.
Supplementary Figure 16: kBET evaluation of batch correction methods.
Supplementary Figure 17: Normalization and cluster annotation.
Supplementary Table 1: Double Application of Antibodies
Supplementary Table 2: Antibody Order Optimization
Supplementary Table 3: Control TMA
Supplementary Table 4: Antibody Order in Replicate TMAs
Supplementary Data 1: Antibodies & Experiments (see Supplementary_Data_1.csv)

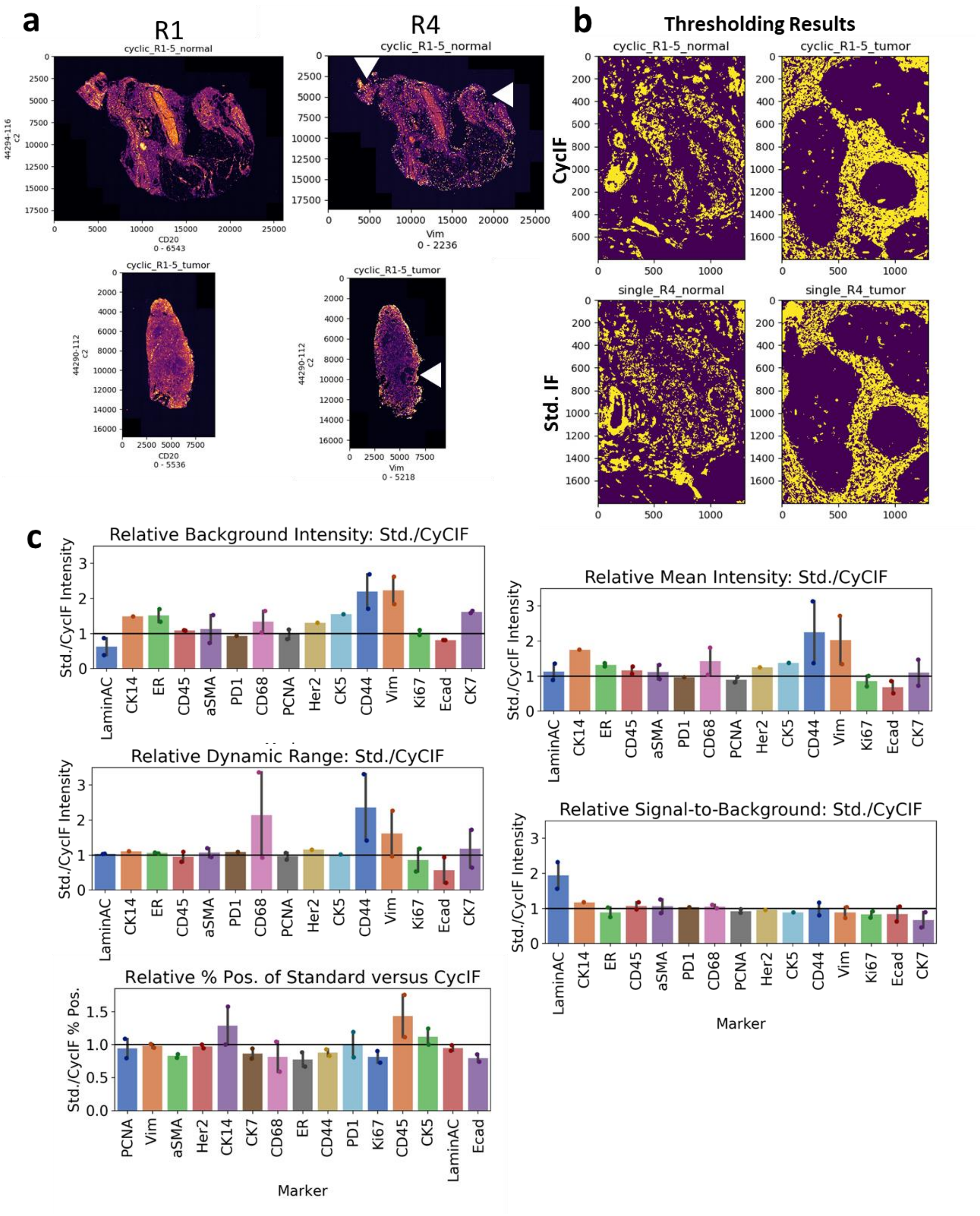


**Supplementary Figure 1: Standard IF versus CycIF in Normal Breast and HER2+ Tumor Tissue.** a. Tissue overview, display range in x-axis label. Round (R) 1 on left, versus R4 on right. Arrowheads show areas of tissue loss. b. Region-of-Interest (ROI) analyzed and mask applied for signal-to-background ratio (SBR) calculation. Yellow=Foreground pixels; background pixels are located 30 pixels (~10 µm) away from foreground pixels. Top panels are CyCIF ROIs and bottom panels are matched standard IF ROIs. c. Intensity, Background, Dynamic Range and SBR and Percent Positive quantification based on threshold in b. (additional markers' thresholds and visualizations here: https://github.com/engjen/cycIF_Validation)

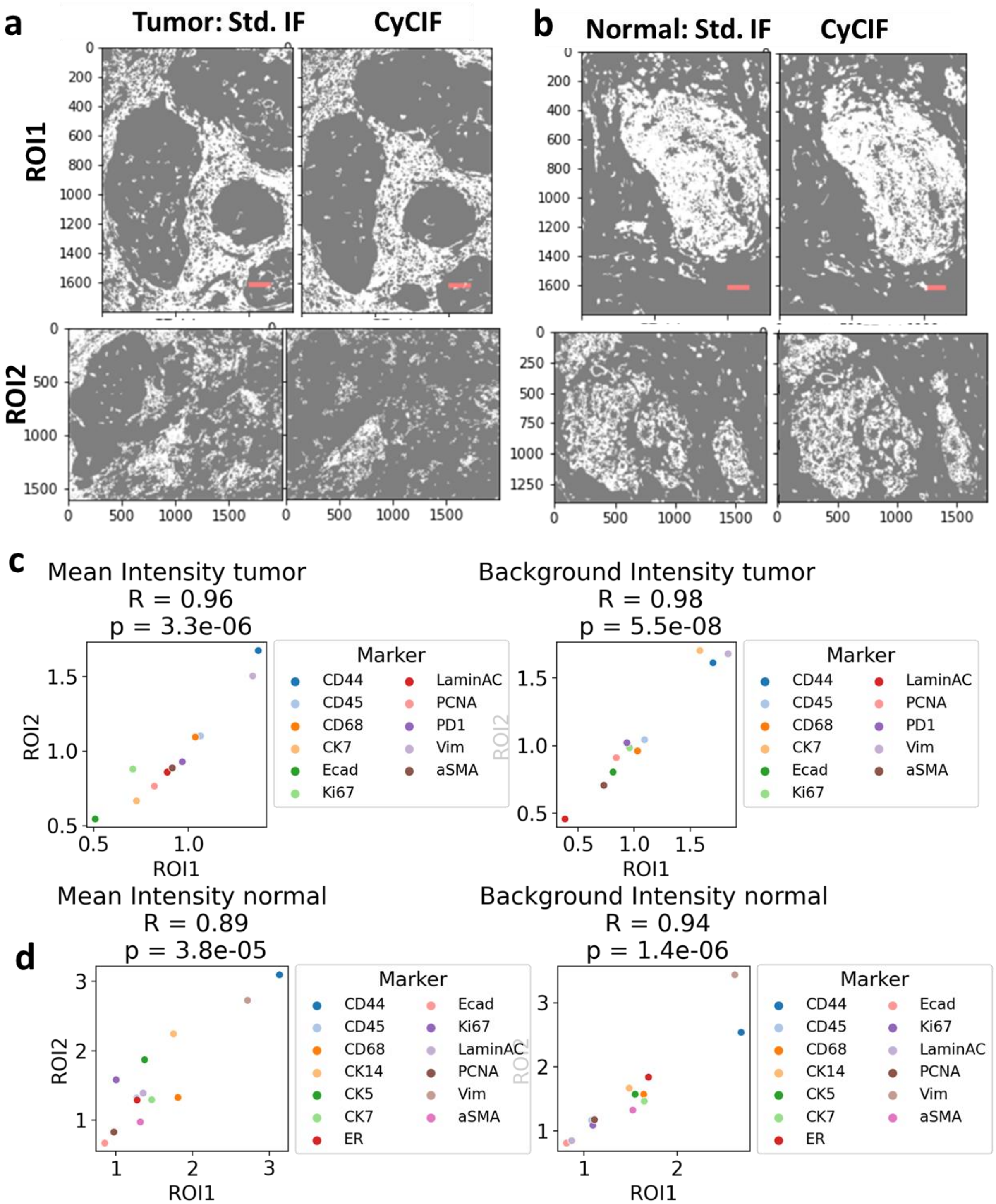


**Supplementary Figure 2: Regional Variation in Threshold Results.** a-b. The same threshold applied to CD44 staining in different ROIs in the tissue, a, tumor tissue and b, normal breast. Top panels show the ROI used for thresholding. Bottom panels show a separate ROI picked from corresponding areas in standard IF and CyCIF slides. Light areas are pixels above threshold, dark areas are pixels below threshold. Comparison of Std. versus CyCIF shows similar pixel patterns in both ROIs. Red scale bar = 50 µm. Axis labels show pixel coordinates; 1 pixel = 0.325 µm. c-d. Correlation of mean positive intensity (left) and background intensity (right) between ROI1 and ROI2. Pearson correlation (R) and p value shown in figure title. c. Correlation of markers expressed in tumor. d. Correlation of markers expressed in normal breast.

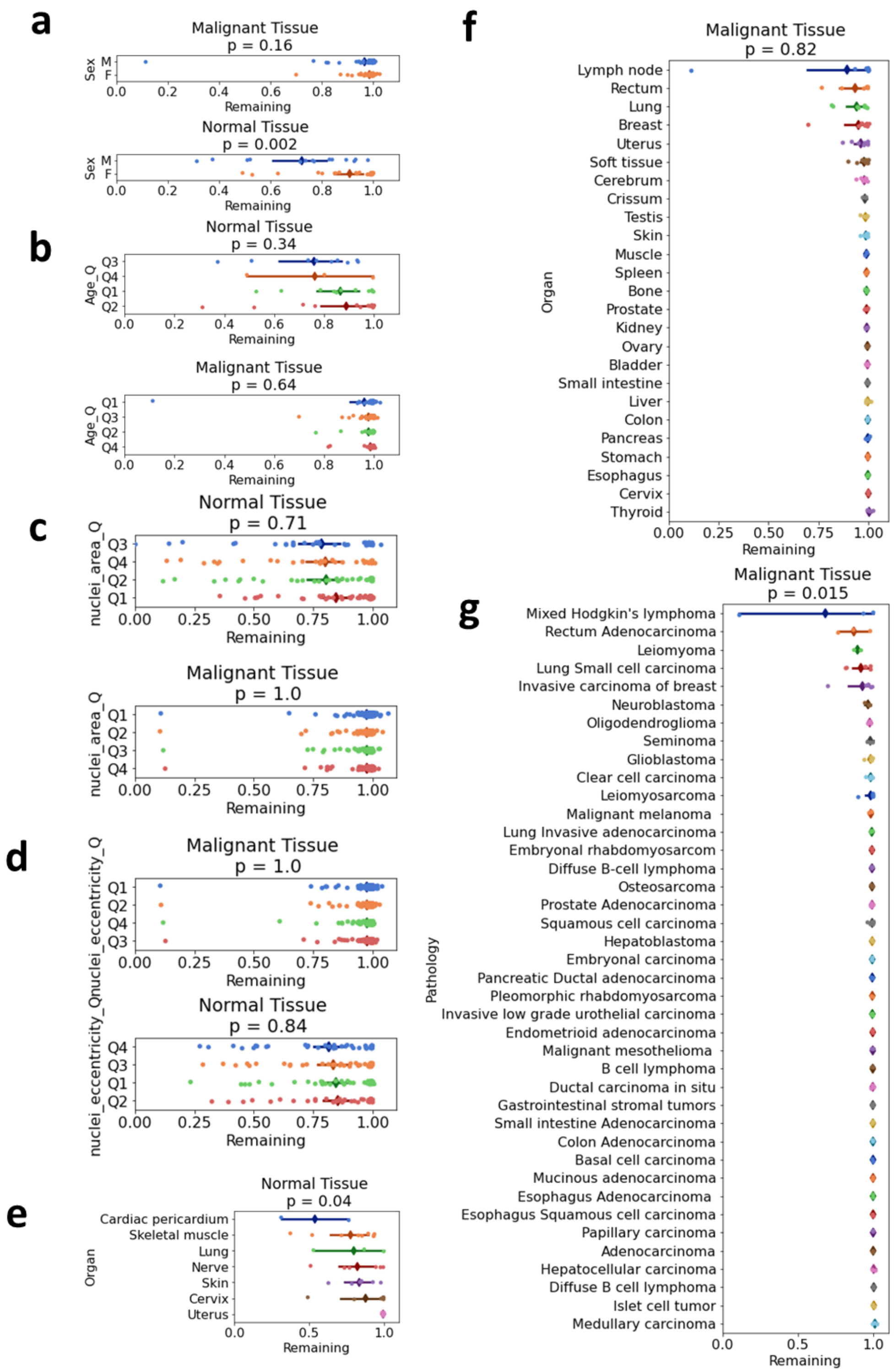


**Supplementary Figure 3: Tissue Retention during CyCIF.** a-g. X-axis shows fraction of cells remaining after 10 rounds of CyCIF. Y-axis shows variables and groups compared. ANOVA was used to assess significant differences; p-value in figure title. n = 156 malignant (top), n = 54 normal (bottom). a. Sex. b. Patient age, by quartile. c. Nuclear area, by quartile. d. Nuclear eccentricity, by quartile. e. Organ, normal tissue. f. Organ, malignant tissue. g. Pathology Diagnosis, malignant tissue.

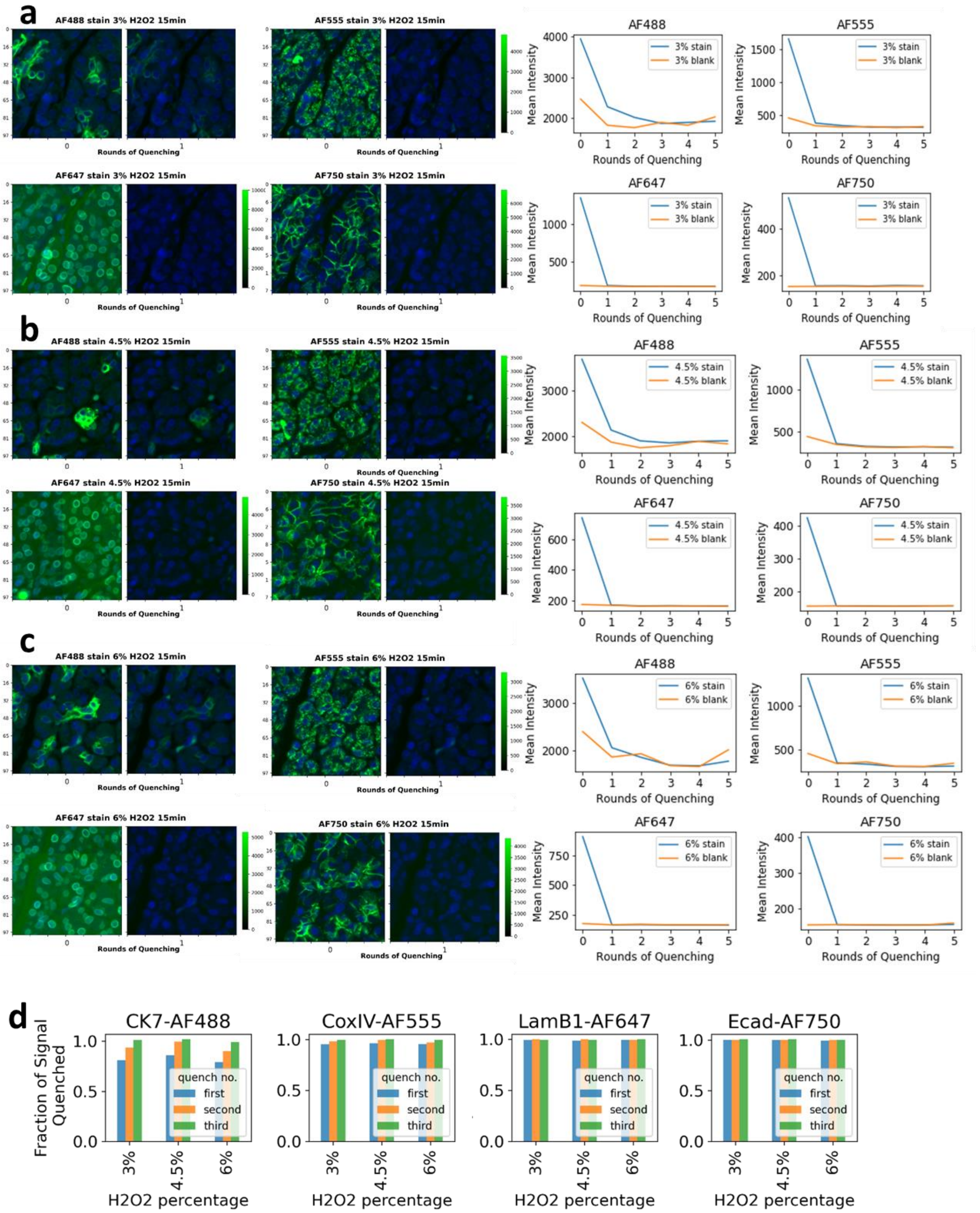


**Supplementary Figure 4: 15-Minute Quenching in Different $H_2O_2$ Concentrations.** a-c. Image panels show staining and after first quench, line plots quantify mean intensity in tissue area over 5 rounds of quenching (blue line). a.3% $H_2O_2$ b. 4.5% $H_2O_2$ c. 6%. $H_2O_2$. Negative controls for all concentrations $H_2O_2$ quantified in a-c (orange line). d. Fraction of signal quenched after first, second and third round of quenching. AF488 takes 3 rounds to quench, AF647 and AF750 only one round.

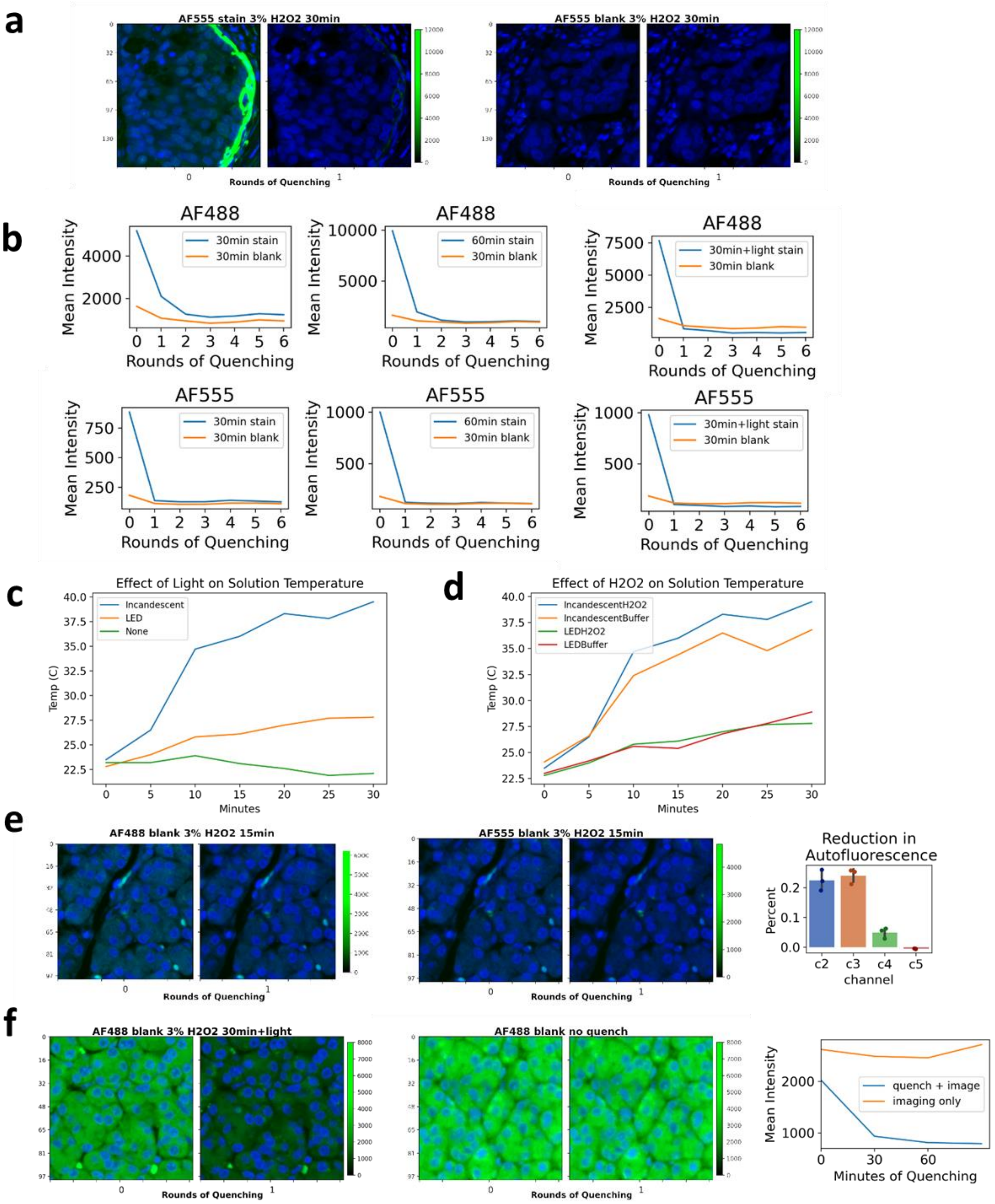


**Supplementary Figure 5. Quenching under Different Time and Light Conditions**. a-b. Quenching performed for 30-minutes, 60 minutes and 30 minutes-plus-light. a. Image panels show AF555 staining and after first quench (left) and blank control (right). b. Line plots quantify mean intensity in tissue area over 6 rounds of quenching, for AF488 (top) and AF555 (bottom), blue line = staining, orange line = blank control. c. Temperature of Quenching Solution under different light conditions. d. Temperature of quenching solution with and without $H_2O_2$, under two light conditions. e. Negative controls in AF488 (left) and AF555 channel (middle), quantified on right, showing reduction in autofluorescence after quenching. f. Blank tissue quenched and imaged (left) versus imaged only (middle) shows quenching is necessary for decrease autofluorescence, seen in comparing blue and orange line in quantification (right).

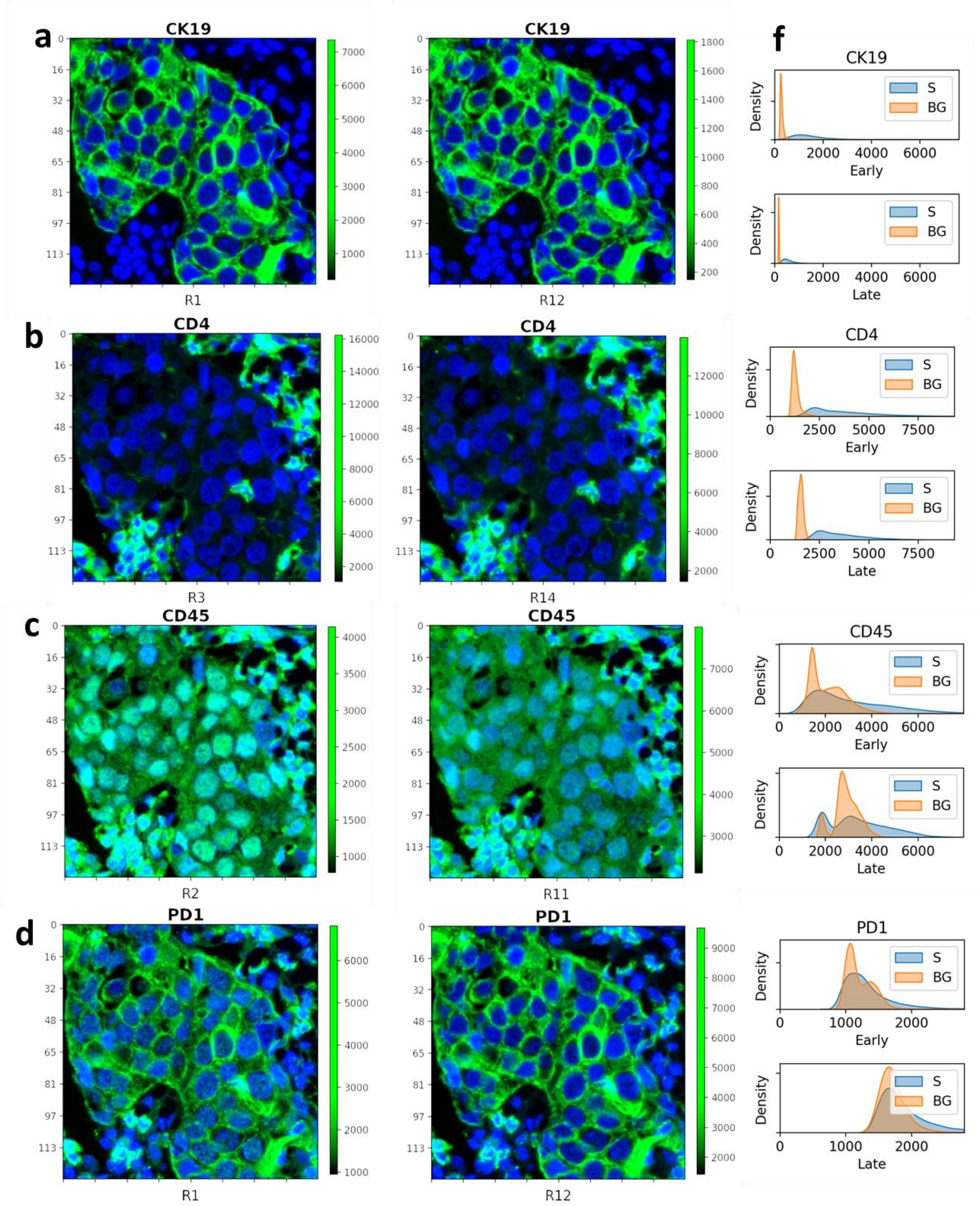


**Supplementary Figure 6: First versus second antibody application to same TMA tissue.** a. CK19 in R1 (first) and again in R12 (second) shows decrease in dynamic range. b. CD4 shows little decrease in dynamic range. c. CD45 shows improvement in non-specific nuclear staining on second application. d. PD1-AF647 has clear bleed trough from CK19-AF750 (matching pattern in a), in both applications. f. Single cell mean intensity distribution of positive signal (S) or background (BG). Cell types were defined by thresholding and gating. (Signal = positive cells for respective marker. Background = Tumor cells for stromal markers, and vice versa. Additional markers' visualizations and figures here: https://github.com/engjen/cycIF_Validation)

a
b
c
d
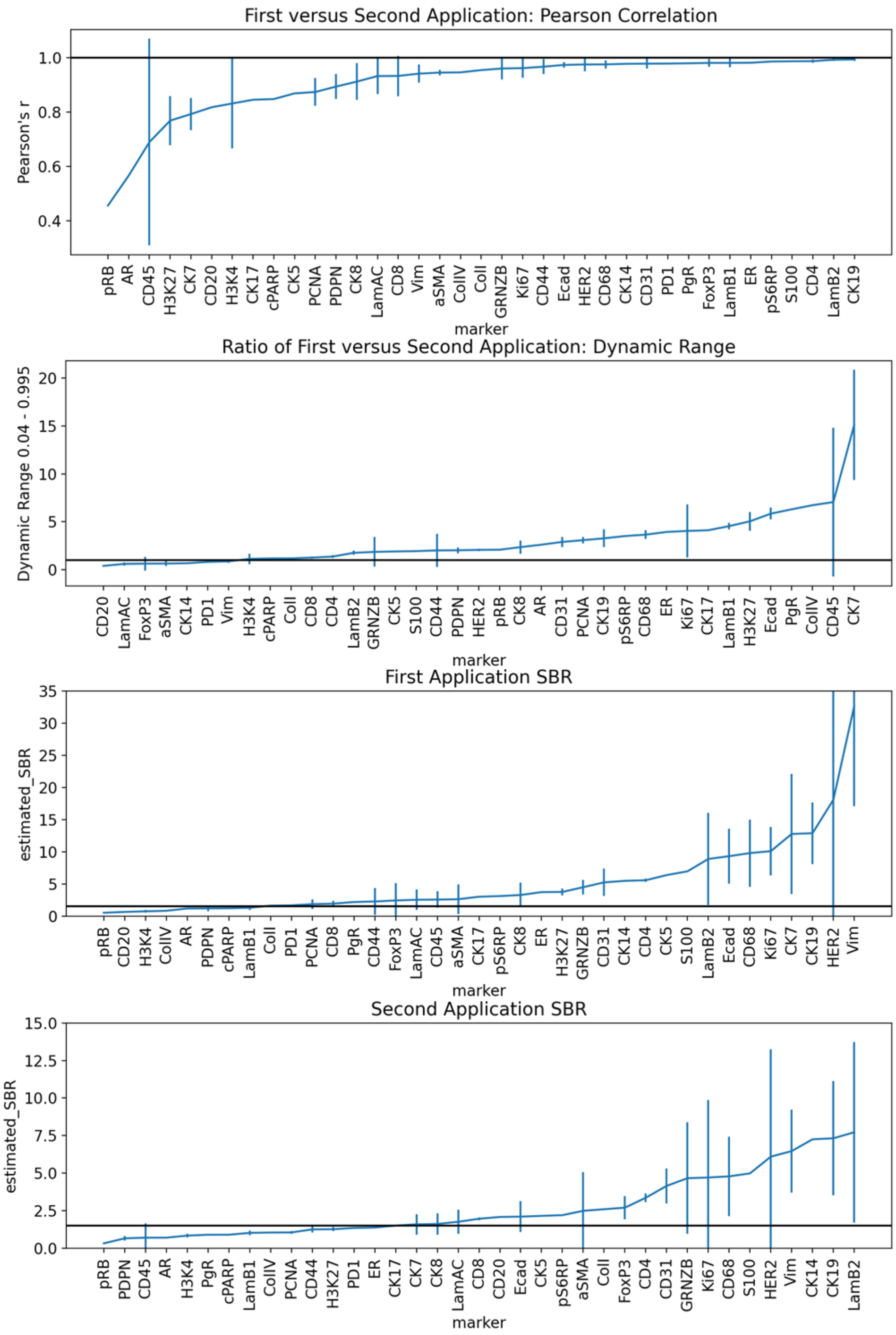


**Supplementary Figure 7: Quantification of first versus second antibody application to same TMA tissue**. a. Pearson Correlation of cells' mean intensity between first application and second application (Error bars S.E.M. n=3). b. Estimated dynamic range (dimmest and brightest 5% of cells, n=3). c-d. Signal to Background ration based on thresholds. 29 of 37 antibodies had SBR above 1.5, while only 23 of 37 second application antibodies had SBR > 1.5 (n=3).

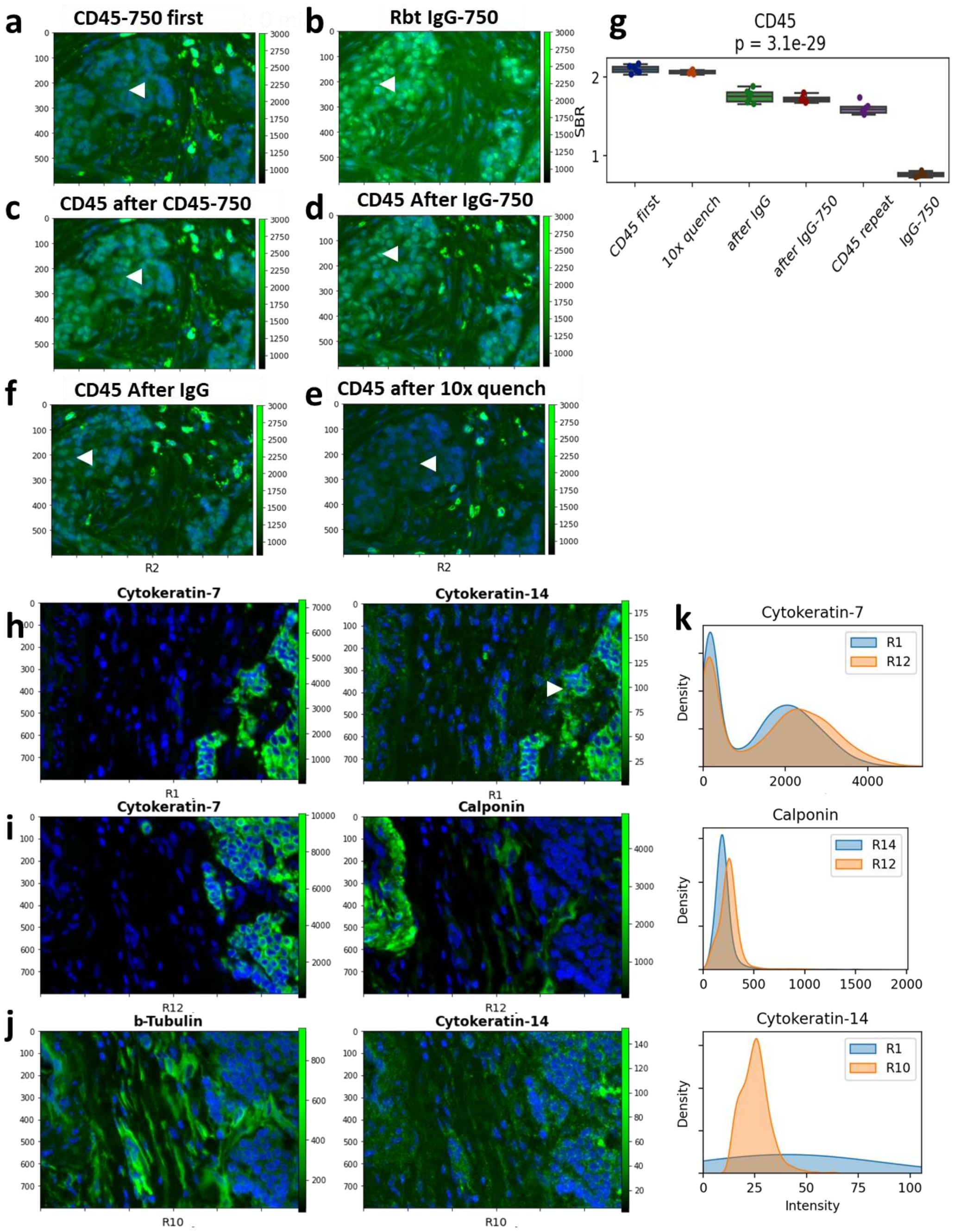


**Supplementary Figure 8: Variables Impacting Round-Effect of Stain Quality and Bleed Through Observed in Miltenyi MACSima prototype Instrument.** a. CD45-750 shows a certain level of non-specific nuclear accumulation (arrowhead). b. Rabbit-IgG antibody labelled with AF750 strongly accumulates in the nucleus or tumor cells (arrowhead). c. Applying CD45-750 again to the same tissue as in (a) increases nuclear background (arrowhead). d. Applying CD45-750 to the same tissue as in (b) increases nuclear background (arrowhead). f. Applying CD45-750 to a tissue previously blocked with unlabeled rabbit-IgG antibody increases nuclear background as well (arrowhead) e. Applying CD45-750 to a tissue previously quenched ten times decreases nuclear background, but also decreases intensity of CD45 staining. g. Signal-to-background quantification of different conditions in a-e. h. Bleed through example in round 1 (R1), from bright Cytokeratin-7-FITC to dim Cytokeratin-14-PE, bleed through shown with arrowhead. i-j. Panel optimization to reduce bleed through. Two bright markers, Cytokeratin-7-FITC and Calponin-PE are placed in R12 (i). Two dim markers, b-Tubulin-FITC and Cytokeratin-14-PE are placed in R10 (j). k. Histograms of single cell mean intensity before and after optimization show Cytokeratin-7 and Calponin are unchanged, and Cytokeratin-14 has reduced bleed through when paired with b-Tubulin in R10.

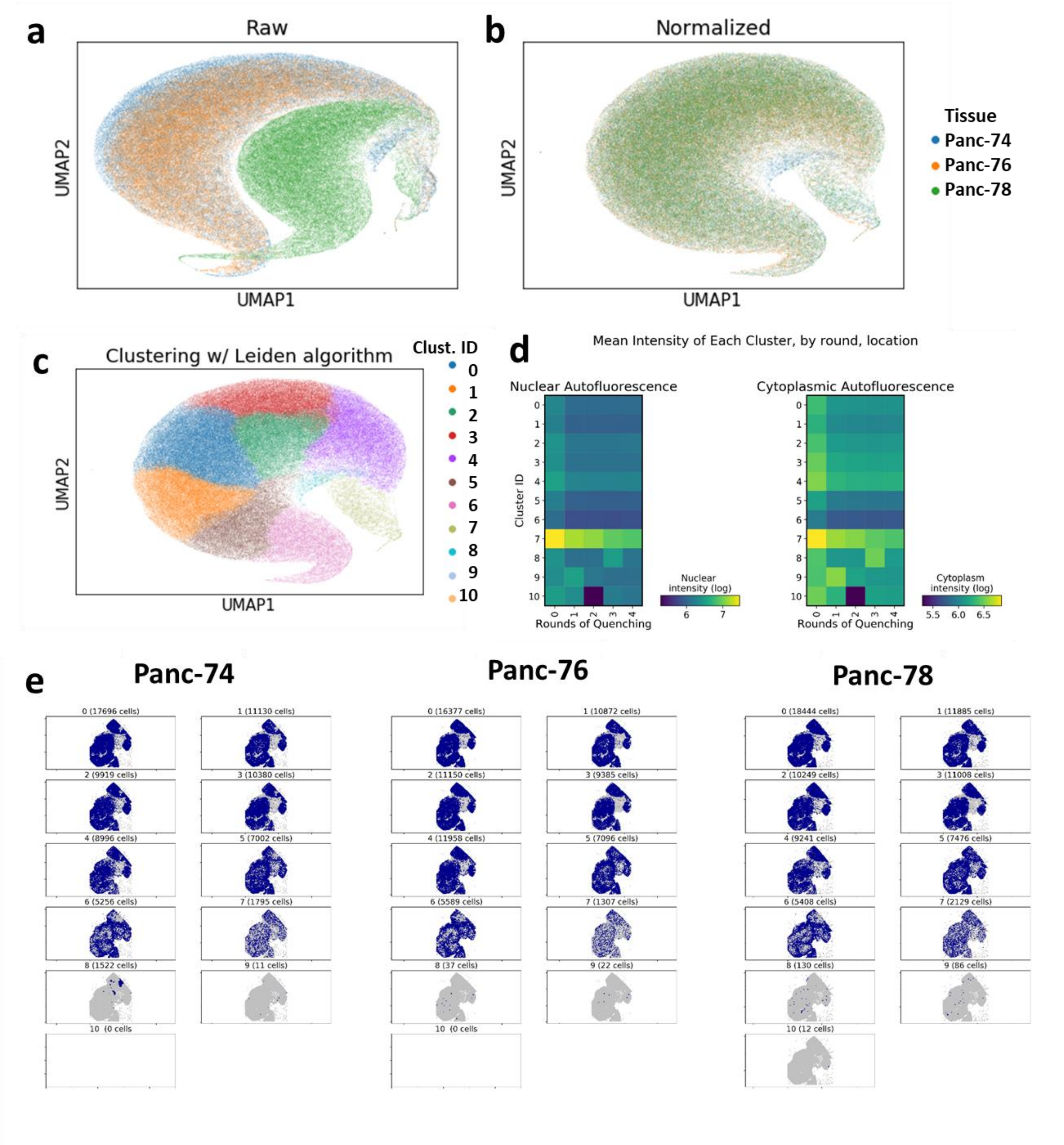


**Supplementary Figure 9. Single Cell Effect of Quenching.** a. UMAP of single-cell autofluorescence in the AF488 channel. b. UMAP of batch-normalized (combat algorithm) single-cell autofluorescence in the AF488 channel. c. Clustering with the Leiden Algorithm yields 10 clusters of similar intensity profiles. d. Mean Autofluorescence in each Leiden Cluster over 0 to 4 15-minute rounds of quenching, separated by subcellular localization. e. Spatial location of cells in each cluster, in each tissue (n=3). Clusters 1-7 show similar localization in all tissues, clusters 8-10 are imaging artifacts.

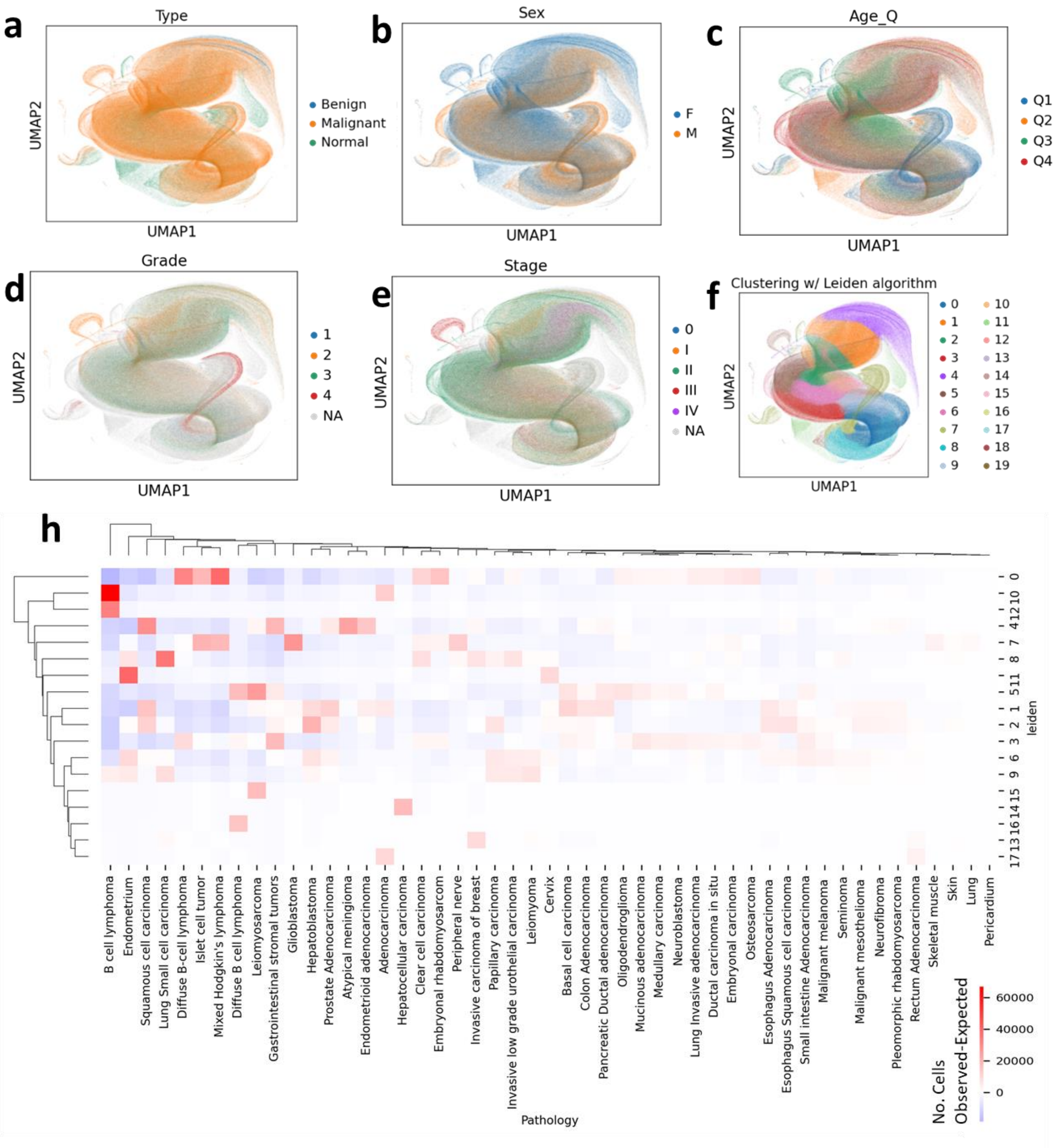


**Supplementary Figure 10. Single Cell Effect of Quenching in 72-core Tissue Microarray.** a-f. UMAP projection based on single-cell autofluorescence in the AF488 channel, colored by: a. Tissue type (normal, benign, malignant). b. Patient sex. c. Patient age, by quartile. d. Tumor grade. e. Tumor stage. f. Leiden cluster ID. h. Heatmap showing observed number of cells per Leiden cluster – expected number of cells per cluster, by pathology diagnosis, y-axis, illustrating different trends by tumor type.

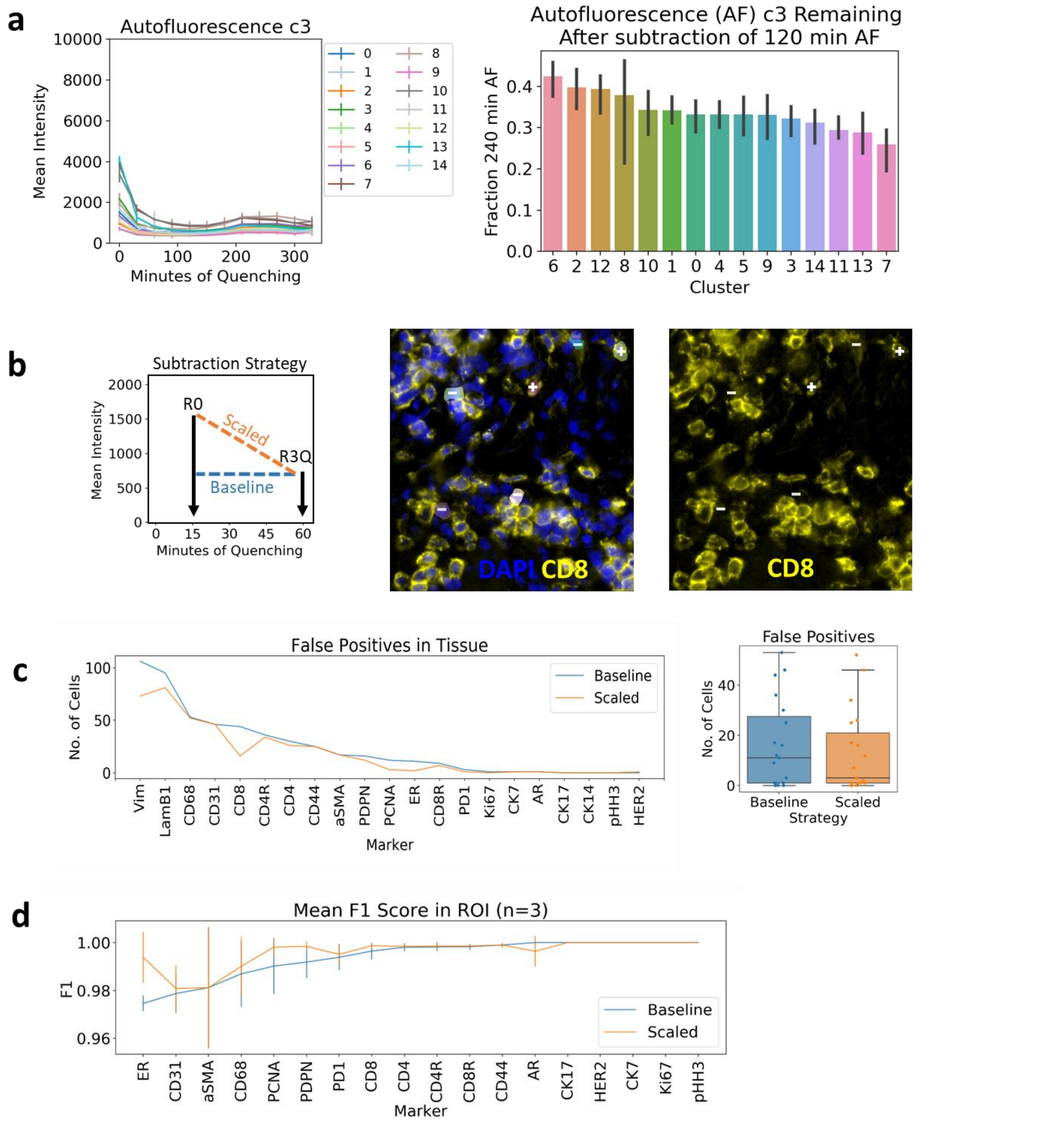


**Supplementary Figure 11. Single Cell Annotation for Evaluation of Autofluorescence Subtraction.** Mean AF555 intensity of cells in each Leiden cluster of 72-core TMA over rounds of quenching (left). Overall, minimum intensity was observed at 120 minutes and maximum intensity at 240 minutes. Subtracting the minimum intensity autofluorescence will avoid over-subtraction while still removing 60 - 70% of autofluorescence at 240 minutes (right). b. Schematic of autofluorescence subtractions strategies, which were applied in (c-d) to a HER2-positive breast cancer tissue (left). Example of CD8 positive/negative annotation in napari image viewer (+ and – overlaid on cell segmentation, center, and CD8 staining, right). c. Number of false positives for each marker (i.e. cells with AF488 autofluorescence > 1024). Boxplots show results of all markers, Mann-Whitney p=0.3. d. Based on single cell annotation in (b), F1 score (true positives)/(true pos + 0.5*(false pos. + false neg.)) of baseline versus scaled subtraction algorithms.

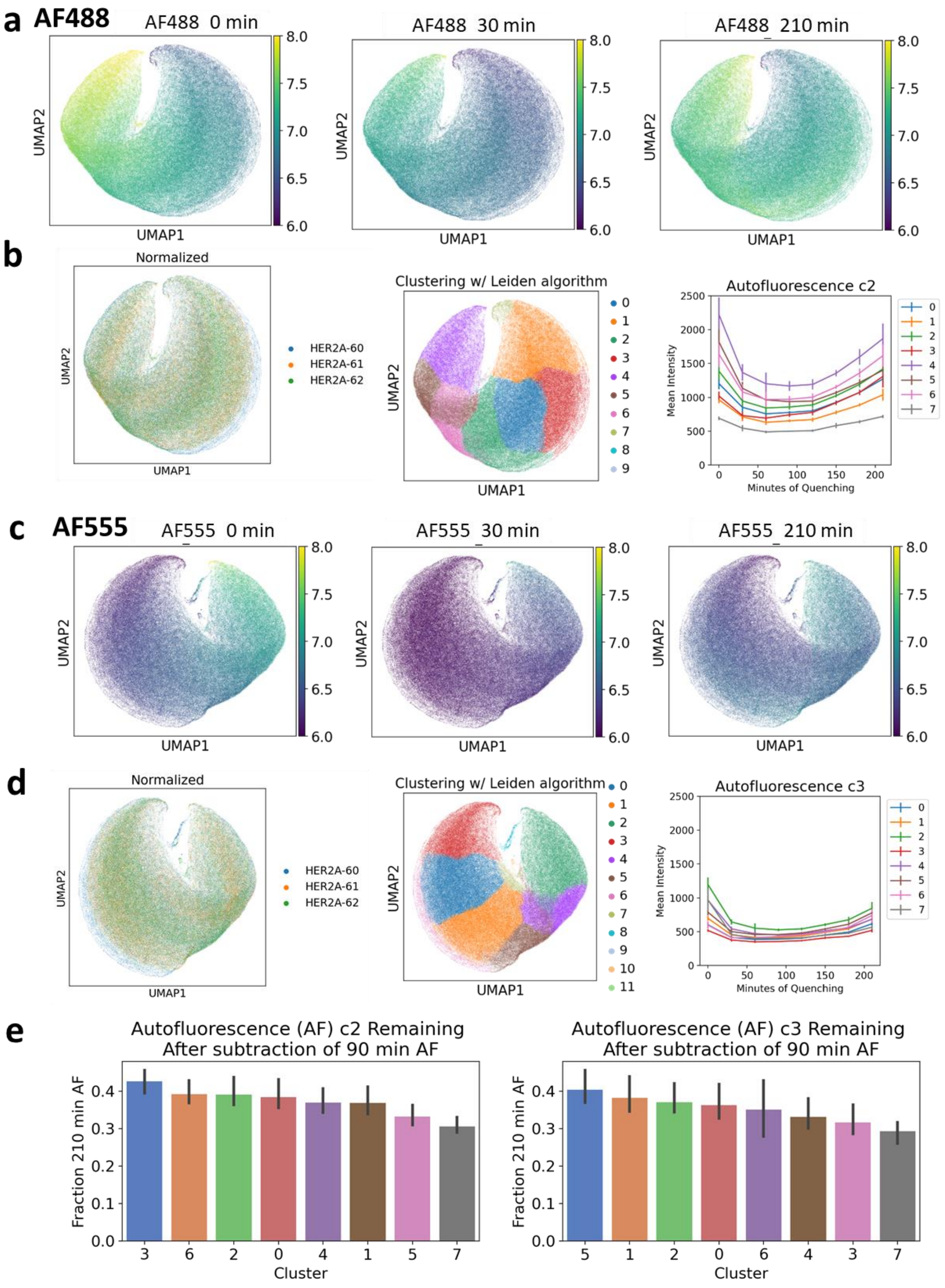


n

**Supplementary Figure 12. Single Cell Effect of Quenching in 11-core HER2+ Tumor Tissue Microarray.** a-b. UMAP projection based on single-cell autofluorescence in the AF488 channel. c-d. UMAP projection based on single-cell autofluorescence in the AF555 channel. a, c. Autofluorescence intensity at 0 (left), 30 (middle) and 210 minutes (right). b, d. UMAP colored by batch (left) - three adjacent sections from TMA were repeatedly quenched and normalized by batch for analysis; UMAP colored by unsupervised clustering results of the Leiden algorithm (middle); Mean autofluorescence intensity of cells in each Leiden cluster over rounds of quenching (right). Overall, minimum intensity was observed at 90 minutes and maximum intensity at 210 minutes. e. Subtracting the minimum intensity autofluorescence will avoid over-subtraction while still removing 60 - 70% of autofluorescence at 210 minutes. (n=3 TMAs x 10 tissues)

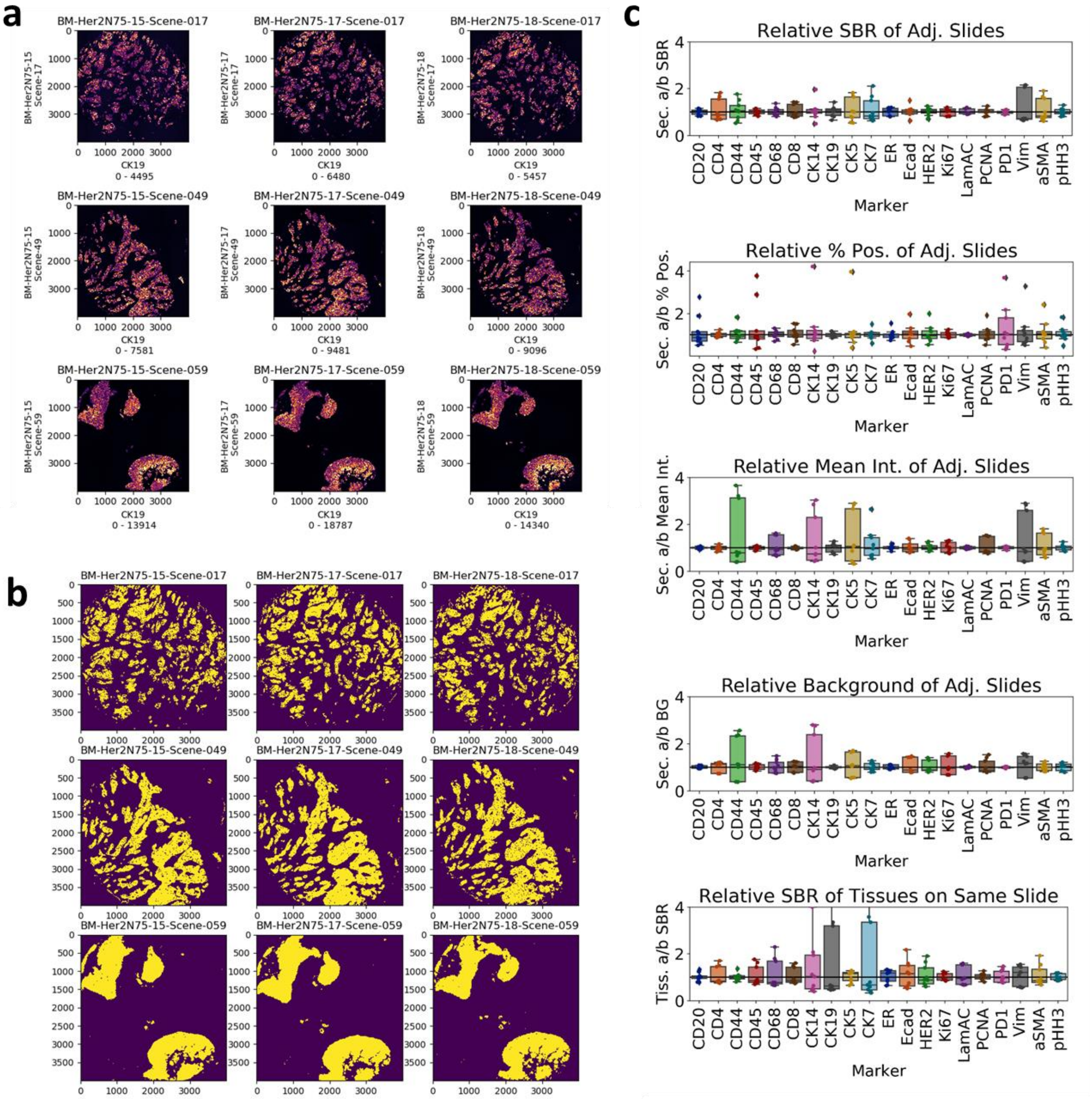


**Supplementary Figure 13: Replicates of Three HER2+ Tumor TMA Cores.** a. Tissue overview, marker and display range in x-axis label. b. Whole core area analyzed; yellow/blue = mask applied for signal-to-background (SBR) calculation. c. SBR, Percent Positive, Intensity, Background, and Tissue-specific SBR quantification, based on threshold in b. (Additional markers' thresholds and visualizations here: https://github.com/engjen/cycIF_Validation)

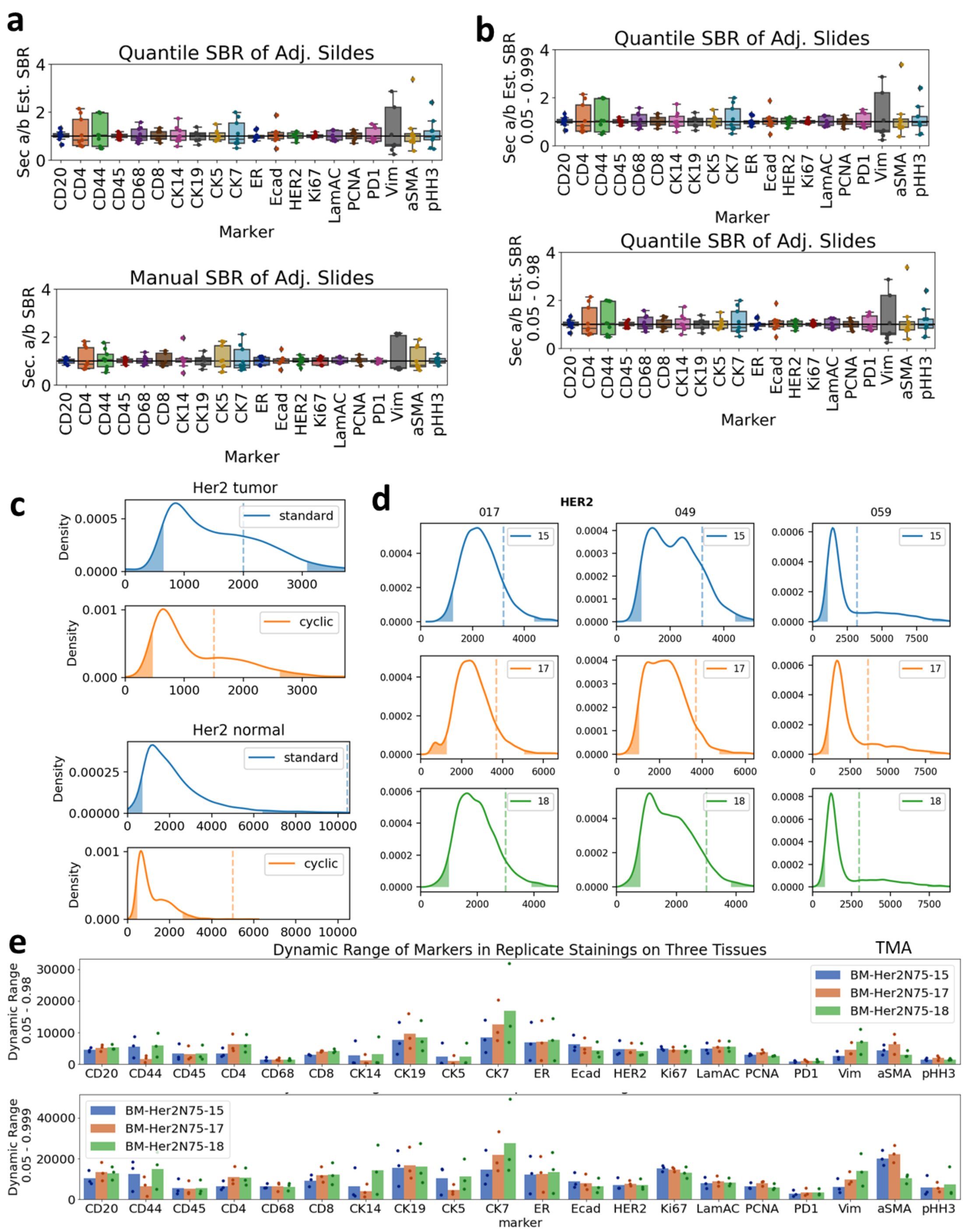


**Supplementary Figure 14: Comparison of Manual Versus Automated SBR and Estimating Dynamic Range with different Quantiles.** Dynamic range of each marker was estimated by the 5th and 95th quantiles of mean intensity. Manual threshold was set to identify positive cells based on staining pattern. a. Replicate cyclic experiments, estimated ("quantile") SBR versus manual. b. We used different quantile ranges to estimate the relative SBR; top 0,05 – 0.999 and bottom, 0.05 to 0.999. The result was not different, indicating the relative quantiles between adjacent slides are stable. c-d. Histograms of single-cell intensity. Dashed line indicates manual threshold, and dotted lines are quantiles for standard versus cyclic (c) and replicate experiments (d). Manual thresholding SBR differs most from estimated SBR in markers that are negative (i.e. HER2 in normal breast, in c). e. Dynamic range of each marker in in three cores in three replicate TMAs. The dynamic range was increased to a greater degree in rare markers, e.g. aSMA and Ki67, when we increase the top quantile form 0.98 (top) to 0.999 (bottom).

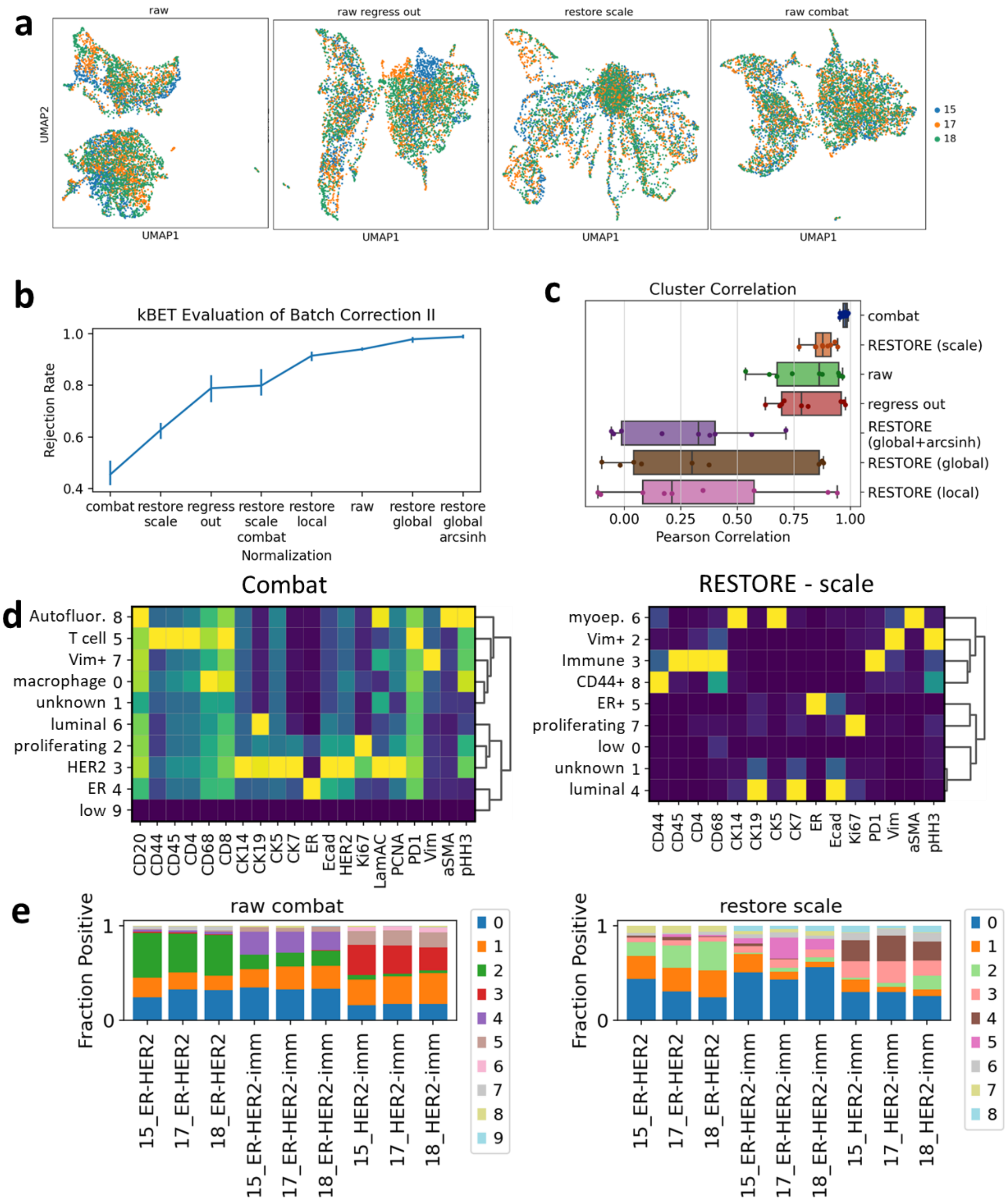


**Supplementary Figure 15: kBET evaluation of batch correction methods in HER2+ breast cancer tissue.** a. Random samples of 1800 cells were drawn from adjacent TMA sections stained with the same cycIF panel. a. UMAP projection of raw or normalized mean intensity, colored by batch. b. kBET rejection rate (lower indicates successful batch correction) was calculated. Error bars S.E.M. (n=3). Several variations of RESTORE algorithm were tested (see methods), in addition to combat and regress out. c. Pearson correlation between replicates' cluster composition, n = 9 (3 cores x 3 replicates). d. Heatmaps of cluster mean intensity on tissue dataset. No autofluorescence (AF) subtraction was performed, and combat showed more influence of AF than RESTORE. e. Cluster composition of replicate TMA cores after combat normalization (left) and RESTORE normalization (right).

**a**

| Transformation | Scaling | Batch regression | Thresholding |
|---|---|---|---|
| raw | standard | regress_out | RESTORE – global |
| log2 | min-max | combat | RESTORE – local |
| arcsinh | max-abs | | RESTORE – scale |
| | robust | | bg-quantile (3rd) |
| | quantile | | |
| | power | | |

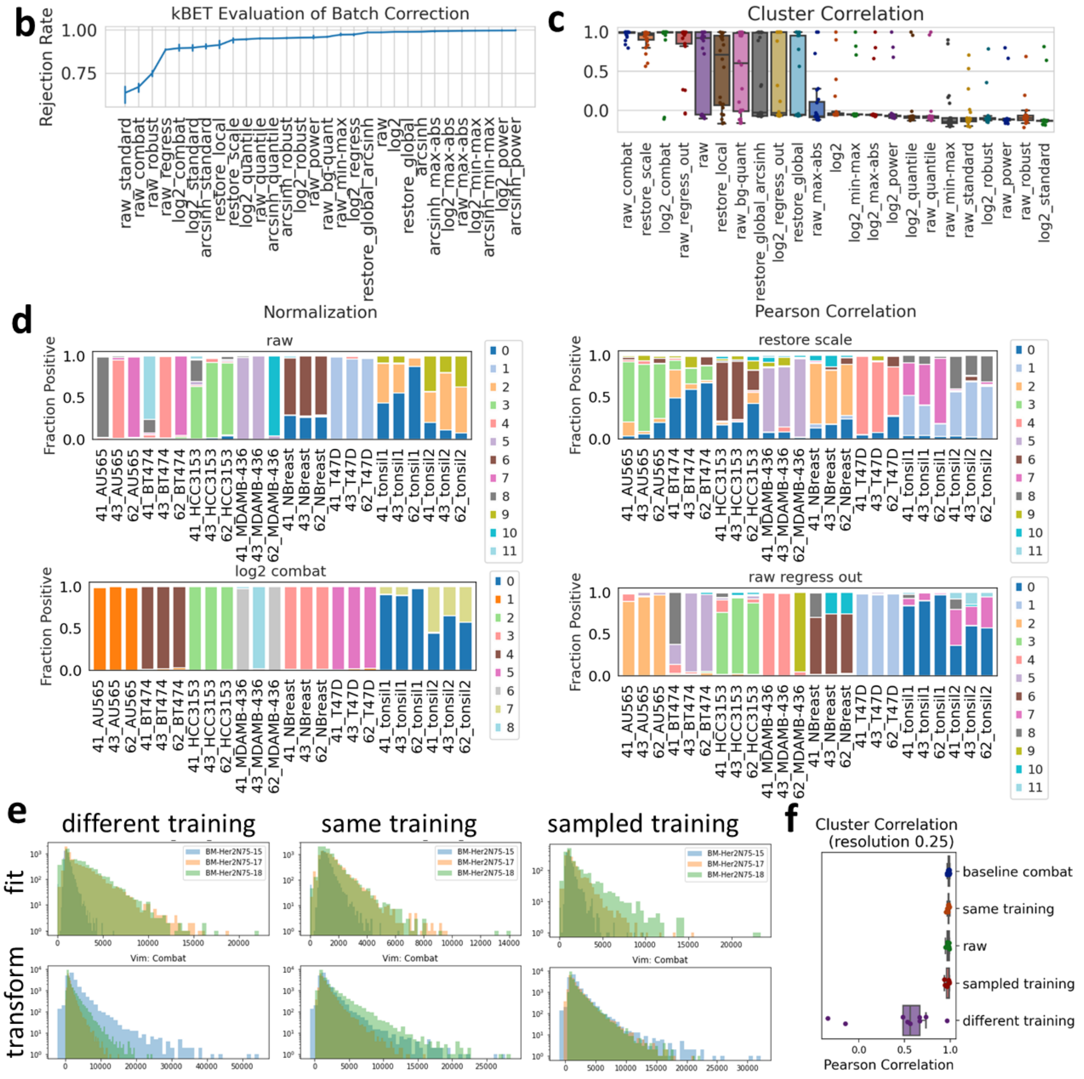


**Supplementary Figure 16: kBET evaluation of batch correction methods.** a. Factors tested for effect on normalization. b. Random samples of 2700 cells were drawn from adjacent TMA sections (from cell lines and normal tissue) stained with the same markers. kBET rejection rate (lower indicates successful batch correction) was calculated. Error bars S.E.M. (n=3). c. UMAP projection colored by batch, of representative batch correction strategies. c. After normalization, unsupervised Leiden clustering was applied (resolution = 0.25) and the Pearson correlation between replicates' cluster composition was calculated; n = 24 (8 cores x 3 replicates). d. Cluster composition of replicate TMA cores, left to right, top to bottom, raw data, RESTORE normalized, log2 transformed and combat normalized, and regress out normalized. The top two clusters per tissue were noted for evaluation of cluster identity. e. Evaluation of effect of training input into combat parameterization and normalization. Combat algorithm was parameterized with a different single tissue within each batch (left), the same single tissue within in batch (center), or a sample of cells from all three tissues in each batch (right). Histograms show intensity distributions used to fit the combat parameters (top) and the resulting normalized data (bottom). f. Under different combat parameterization strategies, unsupervised Leiden clustering was applied (resolution = 0.25) and the Pearson correlation between replicates' cluster composition was calculated.

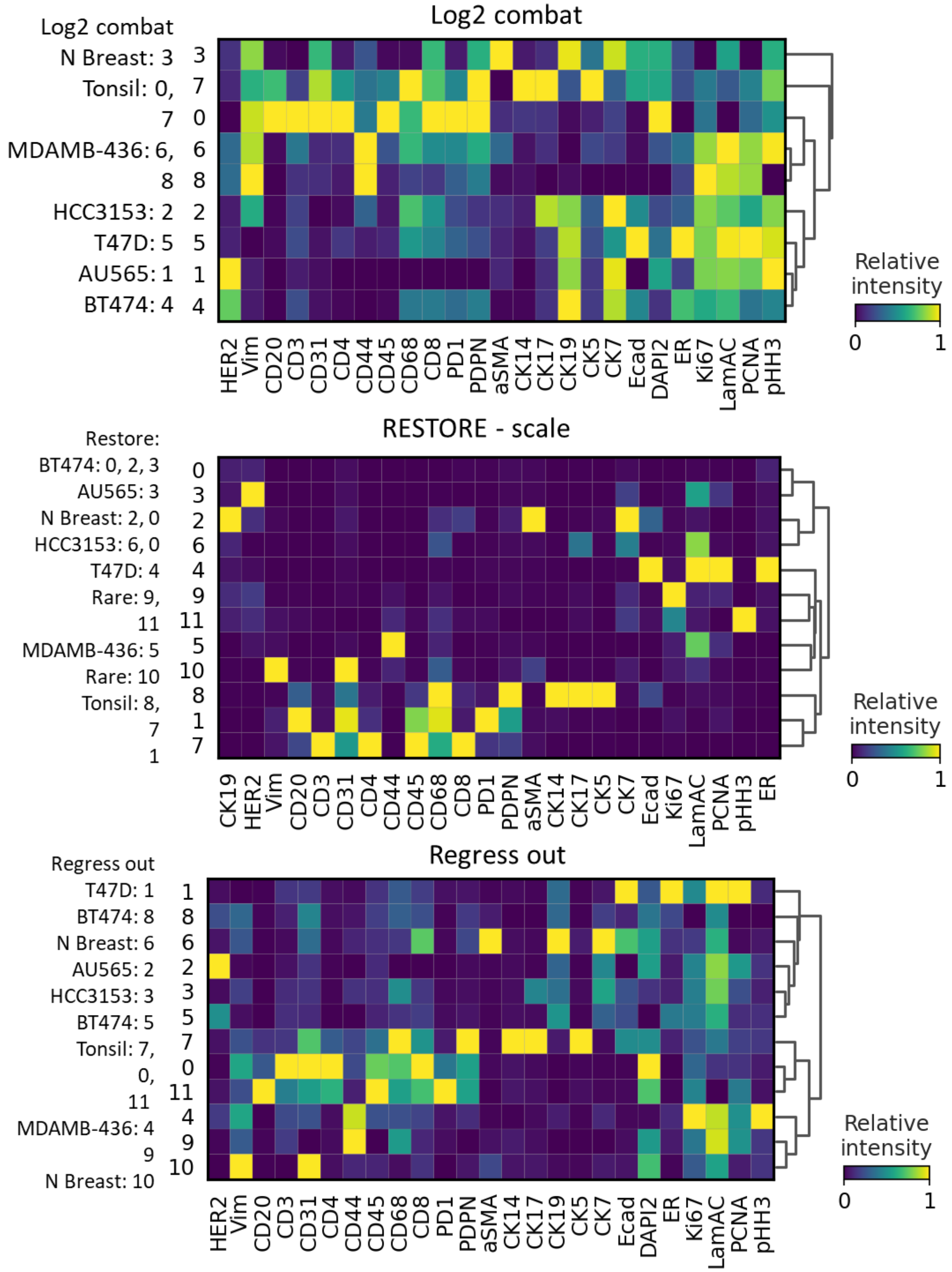


**Supplementary Figure 17: Normalization and cluster annotation.** Unsupervised clustering of normalized data was performed with the Leiden algorithm. For each cell line and normal tissue he top two were noted for evaluation of cluster identity (annotated on left of heat maps). All showed reasonable results, e.g. AU565 should be HER2+, T47D ER+, MDAMB-436 CD44+, BT474 HER2+/ER+, HCC3153 triple negative, tonsil contains immune and basal cytokeratin cell types.

**<u>Tables</u>**

**Supplementary Table 1: Double Application of Antibodies**

| Round | AF488 | AF555 | AF647 | AF750 |
|---|---|---|---|---|
| R1 | PCNA | CD8 | PD1 | CK19 |
| R2 | CK5 | HER2 | ER | CD45 |
| R3 | aSMA | CD68 | CD4 | Ecad |
| R4 | Vim | AR | CD31 | CD44 |
| R5 | CK7 | CK14 | Ki67 | PgR |
| R6 | pHH3 | pS6RP | pERK | CD20 |
| R7 | CK17 | EGFR | pAkt | GRNZB |
| R8 | H3K27 | PDPN | gH2AX | LamAC |
| R9 | CK8 | cPARP | pRB | FoxP3 |
| R10 | LamB1 | H3K4 | ColIV | ColI |
| R11 | S100 | AR* | LamB2 | CD45* |
| R12 | PCNA* | CD8* | PD1* | CK19* |
| R13 | CK5* | HER2* | ER* | CD45** |
| R14 | aSMA* | CD68* | CD4* | Ecad* |
| R15 | Vim* | AR** | CD31* | CD44* |
| R16 | CK7* | CK14* | Ki67* | PgR* |
| R17 | pHH3* | pS6RP* | pERK* | CD20* |
| R18 | CK17* | EGFR* | pAKT* | GRNZB* |
| R19 | H3K27* | PDPN* | gH2AX* | ColI* |
| R20 | CK8* | cPARP* | pRB** | LamAC* |
| R21 | LamB1* | H3K4* | ColIV* | FoxP3* |
| R22 | S100* | CD8** | LamB2* | CD3 |

* second application    ** third application

**Supplementary Table 2: Antibody Order Optimization**

| Marker | Original | Optimized |
|---|---|---|
| aSMA | 3 | 3 |
| CD4 | 3 | 7 |
| CD8 | 1 | 7 |
| CD20 | 6 | 8 |
| CD31 | 4 | 10 |
| CD44 | 4 | 3 |
| CD45 | 2 | 6 |
| CD68 | 3 | 6 |
| CK5 | 2 | 4 |
| CK7 | 5 | 6 |
| CK8 | 9 | 8 |
| CK14 | 5 | 1 |
| CK17 | 7 | 9 |
| CK19 | 1 | 1 |
| ColI | 8 | 9 |
| ColIV | 10 | 8 |
| Ecad | 3 | 2 |
| ER | 2 | 2 |
| FoxP3 | 10 | 5 |
| GRNZB | 7 | 10 |
| HER2 | 2 | 2 |
| LamAC | 11 | 4 |
| LamB2 | 11 | 12 |
| PCNA | 1 | 2 |
| PD1 | 1 | 6 |
| PDPN | 8 | 5 |
| PgR | 5 | 7 |
| pS6RP | 6 | 10 |

**Supplementary Table 3: Control TMA**

| Scene | Cell Line/Tissue | Clinical Sub. | Neve et al. |
|---|---|---|---|
| 10, 11 | AU565 | HER2 + | Luminal |
| 8, 9 | BT474 | ER+/HER2+ | Luminal |
| 2 | HCC1143 | TN | Basal A |
| 3 | HCC3153 | TN | Basal A |
| 12, 13 | MDAMB436 | TN | Basal B |
| 5, 6 | T47D | ER+/PR+ | Luminal |
| 4 | Nor. breast | na | na |
| 1,7 | Tonsil | na | na |

**Supplementary Table 4: Antibody Order in Replicate TMAs**

| Marker | TMA-41 | TMA-43 | TMA-62 |
|---|---|---|---|
| aSMA | 5 | 3 | 4 |
| CD20 | 7 | 8 | 6 |
| CD3 | 6 | 11 | 4 |
| CD31 | 6 | 10 | 6 |
| CD4 | 1 | 7 | 7 |
| CD44 | 3 | 3 | 3 |
| CD45 | 2 | 6 | 5 |
| CD68 | 5 | 6 | 6 |
| CD8 | 1 | 7 | 7 |
| CK14 | 3 | 1 | 1 |
| CK17 | 6 | 9 | 5 |
| CK19 | 1 | 1 | 1 |
| CK5 | 7 | 4 | 2 |
| CK7 | 4 | 6 | 6 |
| DAPI2 | 2 | 2 | 2 |
| Ecad | 5 | 2 | 2 |
| ER | 2 | 2 | 2 |
| HER2 | 2 | 2 | 2 |
| Ki67 | 5 | 1 | 1 |
| LamAC | 4 | 4 | 10 |
| PCNA | 2 | 2 | 3 |
| PD1 | 4 | 6 | 8 |
| PDPN | 6 | 5 | 5 |
| pHH3 | 3 | 5 | 1 |
| Vim | 4 | 7 | 7 |
